# Innate Immune Receptor NLRX1: Potential Modulator of Glioblastoma Pathophysiology

**DOI:** 10.1101/2024.09.19.613932

**Authors:** Durgesh Meena, Divya Shivakumar, Sushmita Rajkhowa, Neermita Bhattacharya, Priya Solanki, Shalini Chhipa, Vikas Janu, Mayank Garg, Jaskaran Singh Gosal, Sushmita Jha

## Abstract

Gliomas are primary brain tumors that develop from glial cells within the central nervous system and are among the deadliest human cancers. Glioblastoma (GBM) is the most malignant form of glioma. NLRX1 is an innate immune pattern recognition receptor that exhibits tumor-suppressive and tumor-promoting effects that may be cancer or cell-type, context-dependent, aided by differences in the microenvironment. Here, we report that NLRX1 is differentially expressed in microglia, astrocytes, GBM cell lines, and glioma patient tissues. siRNA-mediated silencing of *Nlrx1* decreases the ability of the GBM cell line, LN-229, to proliferate and migrate. *Nlrx1*^-/-^ GBM cells exhibit attenuated ability to generate 3D spheroids and enhanced capability to form tunneling nanotubes. Moreover, *Nlrx1^-/-^* GBM cells show decreased expression of autophagy markers, suggesting that NLRX1 plays a role in maintaining autophagy in GBM. In summary, our findings indicate that NLRX1 may modulate GBM pathophysiology by regulating GBM cell proliferation, migration, and metabolism. We believe our understanding of NLRX1 in GBM pathophysiology paves the potential development of GBM-targeting therapeutics that may delay disease progression and/or improve survival.

## Introduction

Gliomas are primary central nervous system (CNS) tumors that develop from glial cells or their progenitors (Louis et al., 2021, 2007). Grade IV glioma, previously known as glioblastoma (GBM), is among the deadliest human cancers. The recently updated WHO classification integrates morphologic and genomic data and classifies GBM into a histologically and genetically defined group consisting of isocitrate-dehydrogenase (IDH)-wild-type diffuse astrocytoma with TERT (Telomerase reverse transcriptase) promoter mutation, or EGFR (Epidermal growth factor receptor) gene amplification, or +7/−10 chromosomal copy number changes (Louis et al., 2021). The current standard of care for the treatment of GBM includes a surgical resection of the tumor followed by treatment with alkylating agent temozolomide and radiation. While these standardized strategies have proved beneficial, they remain essentially palliative as the median survival of GBM patients remains at about 14.6 months (Brown et al., 2016; Chinot et al., 2014; “stupp2009,” n.d.). Glioblastoma is a highly heterogeneous tumor that exhibits astrocytic, oligodendrocytic, neural progenitor, and mesenchymal features (BV and Jolly, 2024; Neftel et al., 2019). One of the leading causes of GBM mortality is the cellular heterogeneity within the GBM tumor microenvironment (TME) (Becker et al., 2021). Patients with GBM possess inter and intratumor heterogeneity (Vollmann-Zwerenz et al., 2020). The existing standard of care leads to GBM recurrence due to the inability to fully eradicate invasive cells and, therefore, results in a poor prognosis (Vollmann-Zwerenz et al., 2020). In the last decade, clinical trials investigating targeted therapies have failed to demonstrate any significant improvement in survival, and innovative new treatments are urgently needed (Brown et al., 2018). Recently, the focus has shifted towards novel strategies that modulate the immune response towards the tumor and the surrounding TME. GBM tumors contain a substantial number of immune cells. Numerous investigations have revealed that 30–50% of the GBM tumor mass comprises microglia/macrophages (Hambardzumyan et al., 2015; Khan et al., 2023). Within the GBM TME, GBM-associated macrophages and microglia produce an immunosuppressive microenvironment by increasing the production of immunosuppressive cytokines (such as Interleukin-6 (IL-6) and Interleukin-10 (IL-10)) and decreasing the expression of immunopermissive cytokines (such as Interleukin-1β (IL-1β)) (Zhang et al., 2011, 2009). Immunotherapy aims to harness the immune system against tumors, with breakthroughs observed in malignant melanoma and haematological malignancies (Brown et al., 2018). However, translating these approaches into therapies for GBM represents a distinct challenge due to the unique TME. Moreover, a transient increase in tumor size can result from inflammatory infiltrates, which within the fixed size of the cranial vault may cause raised intracranial pressure, requiring urgent medical or surgical interventions. Immune-based therapies targeting either tumor-associated macrophages or microglia (TAMs) or the adaptive immunity for glioblastoma are promising, with many clinical trials under progress (Quail et al., 2016; Touat et al., 2017; Weenink et al., 2020). However, recent negative early clinical trials point to the need for a better understanding of tumor-immune cross-talk for patient stratification (Sener et al., 2022).

Inflammation is a protective immune response mediated by innate immune cells to pathogen-associated and danger-associated molecular patterns (PAMPs and DAMPs) as stimuli (Li and Wu, 2021). Pattern recognition receptors (PRRs) recognize these stimuli and activate downstream signaling cascades. The NLR (nucleotide-binding leucine-rich repeat receptors) family of intracellular proteins plays a critical role in innate immunity (Sharma and Jha, 2016). NLRX1 is a member of the NLR family expressed in mitochondria and cytoplasm (Tattoli et al., 2008). The functions of NLRX1 correlate with its subcellular localization. In mitochondria, NLRX1 negatively regulates antiviral signaling through direct interaction with the mitochondrial antiviral-signaling protein (MAVS) on the outer membrane (Allen et al., 2011). This interaction negatively regulates antiviral signaling through negative regulation of Interleukin-6 (IL-6), Interferon-1(IFN-1) production, and possibly NLRP3 inflammasome formation. However, others reported that NLRX1 does not inhibit MAVS signaling (Soares et al., 2013). Some studies have demonstrated that NLRX1 targets the mitochondrial matrix through a functional N-terminal mitochondrial targeting sequence (Arnoult et al., 2009). In the mitochondrial matrix, NLRX1 interacts with UQCRC2 (ubiquinol-cytochrome c reductase core protein 2), a subunit of complex III of the electron transport chain. This interaction induces the generation of reactive oxygen species (ROS), which in turn activates the JNK (c-Jun N-terminal kinase) pathway and results in apoptosis (Tattoli et al., 2008). NLRX1 can be associated with the mitochondrial immune signaling complex (MISC) and TUFM (Mitochondrial Tu translation elongation factor) to promote autophagy (Lei et al., 2012a). In many cancer types, cell and tissue context-specific tumor-suppressive and tumor-promoting functions of NLRX1 have been observed. For example, in colitis-associated carcinogenesis, deletion of *Nlrx1* results in increased expression of tumor necrosis factor (Tnf), epidermal growth factor (Egf), and transforming growth factor beta-1 (Tgfb1), three factors essential for wound healing, and increased epithelial proliferation following injury triggered by dextran sodium sulfate (Tattoli et al., 2016). Similar results were also observed in primary intestinal organoids. Deletion of *Nlrx1* in primary intestinal organoids results in exacerbated proliferation and expression of intestinal stem cell markers upon TNF stimulation. This hyper-proliferation response was associated with increased activation of Protein kinase B (Akt) and Nuclear factor-kappa B (NF-kB) pathways in response to TNF stimulation. Further studies supported the tumor-suppressing role of NLRX1 in pancreatic and primary breast cancer (Nagai-Singer et al., 2023; Singh et al., 2015). In contrast to these studies, the tumor-promoting role of NLRX1 was found in basal-like and metastatic breast carcinoma (Singh et al., 2019). In the triple-negative breast cancer cell line, MDA-MB-231, in the presence of TNF-α, loss of *Nlrx1* is associated with altered oxidative phosphorylation system (OxPhos) activity, impaired lysosomal function, and mitophagy. Further depletion of *Nlrx1* decreased OxPhos-dependent cell proliferation and migration ability of triple-negative breast cancer cells in the presence of TNF-α. Therefore, the impact of NLRX1 on cancer is likely to vary depending on the specific type of cancer or cell type, influenced by variations in the microenvironment. While several studies have shown the tumor-promoting and suppressing role of NLRX1 in different cancer types, the role of NLRX1 in regulating GBM progression has remained unexplored. This study, therefore, aims to understand differential NLRX1 signaling in GBM and how it regulates GBM pathophysiology, leading to the identification of novel signaling pathways that, in turn, may help create better GBM therapy approaches and improve overall survival.

## Results

### 1. NLRX1 is differentially expressed across GBM and astrocyte cell lines

NLRX1 is a mitochondrial innate immune protein, and it is expressed in various cell lines, including breast cancer cell lines (ZR-55-1, MCF-7, BT-474, T47D, MDA-MB-231, HBL100), embryonic kidney cells (HEK293), cervical cancer cells (HeLa), mouse embryonic fibroblast cells (MEFs), and mouse bone-marrow derived-macrophages. (Singh et al., 2019, 2015; Soares et al., 2014; Zhang et al., 2019). The expression level of NLRX1 varies from cell to cell. NLRX1 expression has not been analyzed in GBM cell lines. We have examined the expression of NLRX1 in various GBM cell lines, including LN-18, LN-229, and U87-MG, as well as in the astrocyte cell line SVG. The endogenous NLRX1 exhibits a punctate distribution in the cytoplasm. Nevertheless, the expression pattern of NLRX1 exhibits variability among different cells, as depicted in Figure 1A. NLRX1 shows elevated expression levels at membrane protrusions in LN-229 cells and is predominantly localized around the nucleus in LN-18 cells. NLRX1 exhibits widespread expression in both SVG and U87-MG cells, encompassing the whole cell, including the nucleus and cytoplasm. The variation in expression pattern may be attributed to distinct signaling pathways that various cell lines adhere to. The LN-229 and LN-18 cells display mutant p53 (TP53) and perhaps homozygous deletions in the p16 and p14ARF tumor suppressor genes. The U87-MG cell line is characterized by a hypodiploid state, meaning it has a chromosome number below the normal diploid level. Specifically, it has a modal chromosome number of 44, observed in 48% of the cells. Additionally, to quantify the expression of NLRX1 in different types of cells, western blot analyses were performed utilizing protein lysates from GBM and astrocyte cell lines (Fig 1B). The densitometry analysis indicates minimal variation in NLRX1 total baseline protein expression across normal and GBM cell types (Fig 1C).

**Figure 1.**
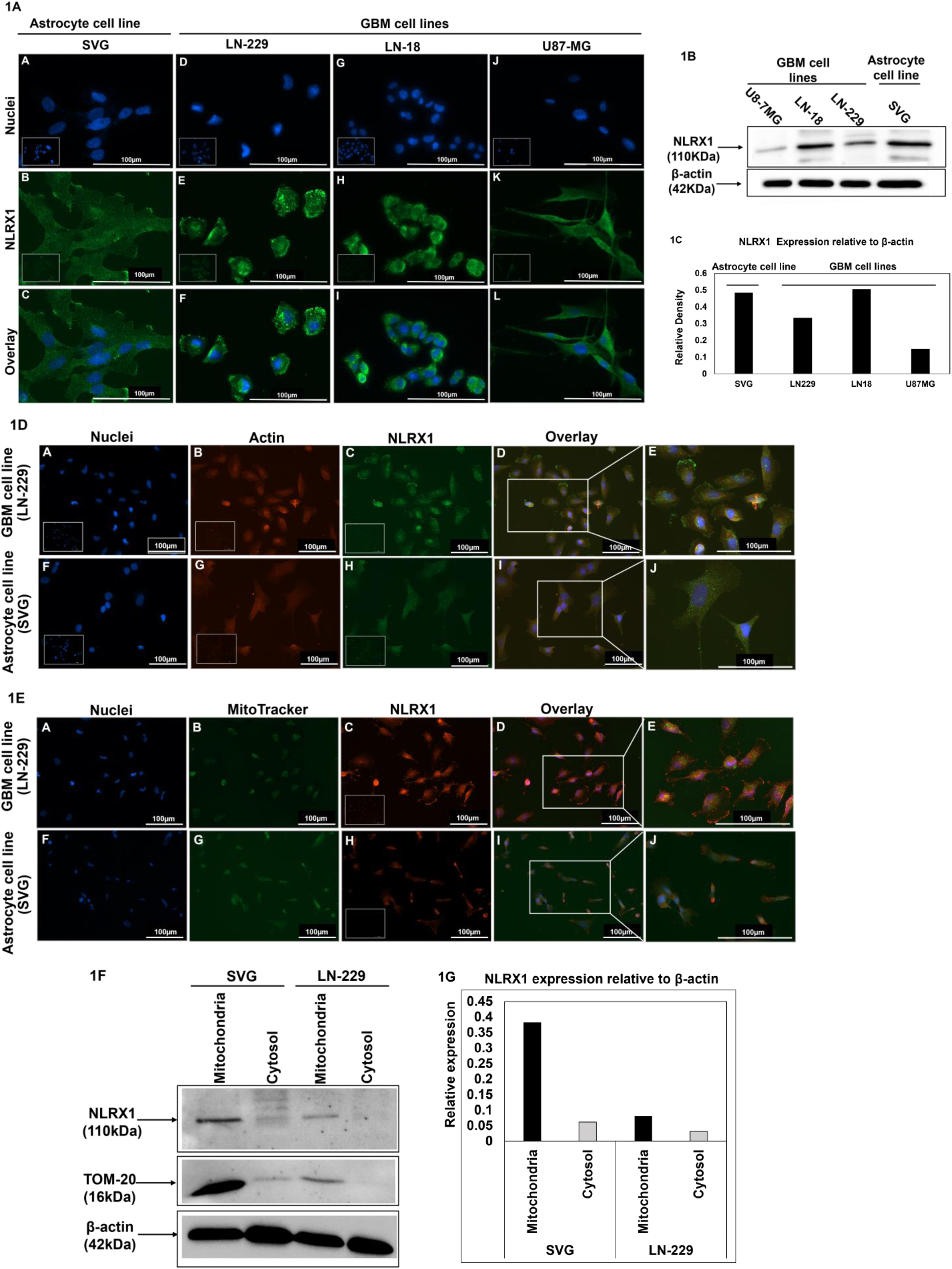
NLRX1 is differentially expressed across GBM and astrocyte cell lines: A. To check the NLRX1 expression across various GBM and astrocyte cell lines, cells were stained with anti-NLRX1 antibody (green) and DAPI (nuclei, blue). At least 7 frames were imaged per well of the two well chamber slides. Images are representative of all frames. Inset represents primary antibody control. Scale bar,100 µm. B. NLRX1 expression is quantified across various cell types using western blot. β-actin is used as a loading control. 20µg protein sample is loaded in each well. Images are representative of 3 experiments. C. The western blot data was quantified using densitometry. The graph represents data from one representative experiment. D. GBM cell line (LN-229) and astrocyte cells (SVG) were stained with anti-actin antibody (red), anti-NLRX1 antibody (green), and DAPI (nuclei, blue). At least 7 frames were imaged per well of the two well chamber slides. Images are representative of all frames. Inset represents primary antibody control. Scale bar, 100 µm. E. GBM (LN-229) and astrocyte cell line (SVG) were stained with MitoTracker (green), anti-NLRX1 antibody (red), and DAPI (nuclei, blue). At least 7 frames were imaged per well of the two well chamber slides. Images are representative of all frames. Inset represents primary antibody control. Scale bar, 100 µm. F. Subcellular NLRX1 expression in mitochondria and cytosol of GBM (LN-229) and astrocyte cell line (SVG) was quantified using a western blot. β-actin is used as a loading control. 10µg protein sample is loaded in each well. Images are representative of 2 experiments. G. The western blot data was quantified using densitometry. The graph represents data from one representative experiment.

Previous studies have demonstrated that NLRX1 is expressed in the cytoplasm, mitochondria, and cell membranes (Moore et al., 2008). However, the subcellular expression pattern of NLRX1 is context and TME-dependent. To examine the subcellular expression pattern of NLRX1 within an un-stimulated GBM (LN-229) and astrocyte cell line (SVG), colocalized staining of NLRX1 with cytoskeleton protein actin, and MitoTracker (a dye used to stain live mitochondria) was performed (Fig 1D, 1E). The merged fluorescent image in Figure 1D displays the co-localization of NLRX1 expression with actin staining, indicating the presence of NLRX1 in the cytoskeleton components of both GBM and astrocyte cell lines. Further, as mitochondria are crucial organelles that are carried within the cell via the cytoskeleton, and the previous studies report the presence of NLRX1 in mitochondria, we examined the expression of NLRX1 in mitochondria of both GBM and astrocyte cell lines. Figure 1E unambiguously demonstrates the expression of NLRX1 in the mitochondria of GBM and astrocyte cell lines. Further, to quantify the NLRX1 expression in various subcellular compartments, the mitochondria and cytosolic fractions of the GBM cell line, LN-229, and astrocyte cell line, SVG were isolated and subjected to a western blot analysis for NLRX1 (Fig 1F). By conducting a densitometry analysis of NLRX1 expression, it has been revealed that the expression of NLRX1 is greater in the mitochondrial fraction compared to the cytosolic fraction in both GBM and astrocyte cell lines (Fig 1G).

### 2. NLRX1 is differentially expressed across diverse glioma patients

GBM is a heterogeneous tumor that consists of a variety of cells, including malignant cells, immune cells, vascular cells, stem cells, and glial cells in its TME. Macrophages and microglia make up approximately 30-40% of all cells in GBM (Hambardzumyan et al., 2015). GBM cells release molecules, such as SDF-1 (stromal-derived factor-1), CCL2 (chemokine [C-C motif] ligand 2), CCL7, and CX3CL1, which aid in attracting macrophages and microglia to the GBM TME (Held-Feindt et al., 2010; Kioi et al., 2010; Okada et al., 2009; Zhang et al., 2012). GBM cells also secrete hepatocyte growth factors, that support the survival of macrophages and microglia (Alghamri et al., 2021; Alterman and Stanley, 1994; Badie et al., n.d.). Microglia in GBM TME promotes tumor cell invasion by producing epidermal growth factor (Coniglio et al., 2012). Astrocytes, a primary constituent of the GBM microenvironment, transform into reactive astrocytes (Zhang et al., 2016). Reactive astrocytes in the TME of GBM engage with microglia, leading to the promotion of an immunosuppressive microenvironment by releasing cytokines such as transforming growth factor beta (TGFβ), interleukin-10 (IL10), and granulocyte colony-stimulating factor (G-CSF) (Henrik Heiland et al., 2019). Therefore, the tumor-immune crosstalk establishes an immunosuppressive network and results in immune evasion (Croci et al., 2007).

In order to assess the expression of NLRX1 in innate immune cell populations in glioma and normal brain tissues, paraffin-embedded patient tissue sections were stained for NLRX1 with microglia and astrocytes (Fig 2A, 2C). For the staining of microglia, RCA (Ricinus Communis Agglutin, a lectin that binds to β-D-galactose residues and labels microglia) was used. For the staining of astrocytes, anti-GFAP (Glial Fibrillary Acidic Protein) antibody was used. The overlay images in Figures 2A and 2C demonstrate the co-localization of GFAP and RCA immunofluorescence with anti-NLRX1 immunofluorescence, confirming the presence of NLRX1 in microglia and astrocytes of both glioma and normal brain tissues. Immunopositive NLRX1-expressing astrocytes and microglia cells with an observable DAPI-stained nucleus were counted blindly twice. Cell counts are averages of at least seven fields (Fig 2B, 2D). The abundance of NLRX1-expressing astrocytes and microglia is greater in normal brain tissue than in glioma tissues (p < 0.005). We also assessed the expression of NLRX1 in primary astrocytes and microglia. This is because the protein expression may vary between primary cells and cell lines or fixed tissue samples. Primary cells derived from freshly obtained glioma tissues were extracted and subjected to staining for astrocytes, microglia, and NLRX1 (Fig 2E, 2F). The overlay images in Fig 2E and 2F demonstrate the co-localization of NLRX1 expression with astrocyte (GFAP) and microglia (RCA) immunofluorescence. This result provides additional evidence of the presence of NLRX1 in primary astrocytes and microglia. Following an examination of the baseline expression of NLRX1 in various cell lines (GBM and astrocytic), innate immune cells (microglia and astrocytes) from fixed patient tissues, as well as glioma patient tissue-derived primary cells, the expression of NLRX1 is quantified across diverse grades of glioma patient tissues (Fig 2G, 2H). Furthermore, the expression of GFAP was quantified alongside NLRX1 to obtain an estimate of the abundance of astrocytes in relation to NLRX1 (Fig 2G, 2I). Adjacent normal tissue from the same patient was used as control. The western blot data demonstrates the varied NLRX1 and GFAP expression among different patients, as well as between the normal and tumor tissues of each glioma patient. These findings corroborate the existing research on the heterogeneity of GBM. Glioblastoma was first described as glioblastoma “multiforme” by Bailey and Cushing in 1922 because of the highly heterogeneous gross presentation (Bianco et al., 2017). The TME also varies in the same tumor as the genes related to oncogenic signaling, proliferation, complement response, immune response, and hypoxia signaling pathways vary from cell to cell in the same tumor. This heterogenous TME of GBM is one of the reasons for GBM recalcitrance, as a single therapy may be effective for one population of cells or resistant for another cell population because of differential oncogenic signaling (Qazi et al., 2017). Reinhartz and group showed that single-cell subclones derived from the same tumor display marked heterogeneity in response to drug treatments (Reinartz et al., 2017). Our data further supports the inter-patient heterogeneity of glioma tumors at gene and molecular profile levels. Table 1 lists the details of the patients used for the study.

**Figure 2.**
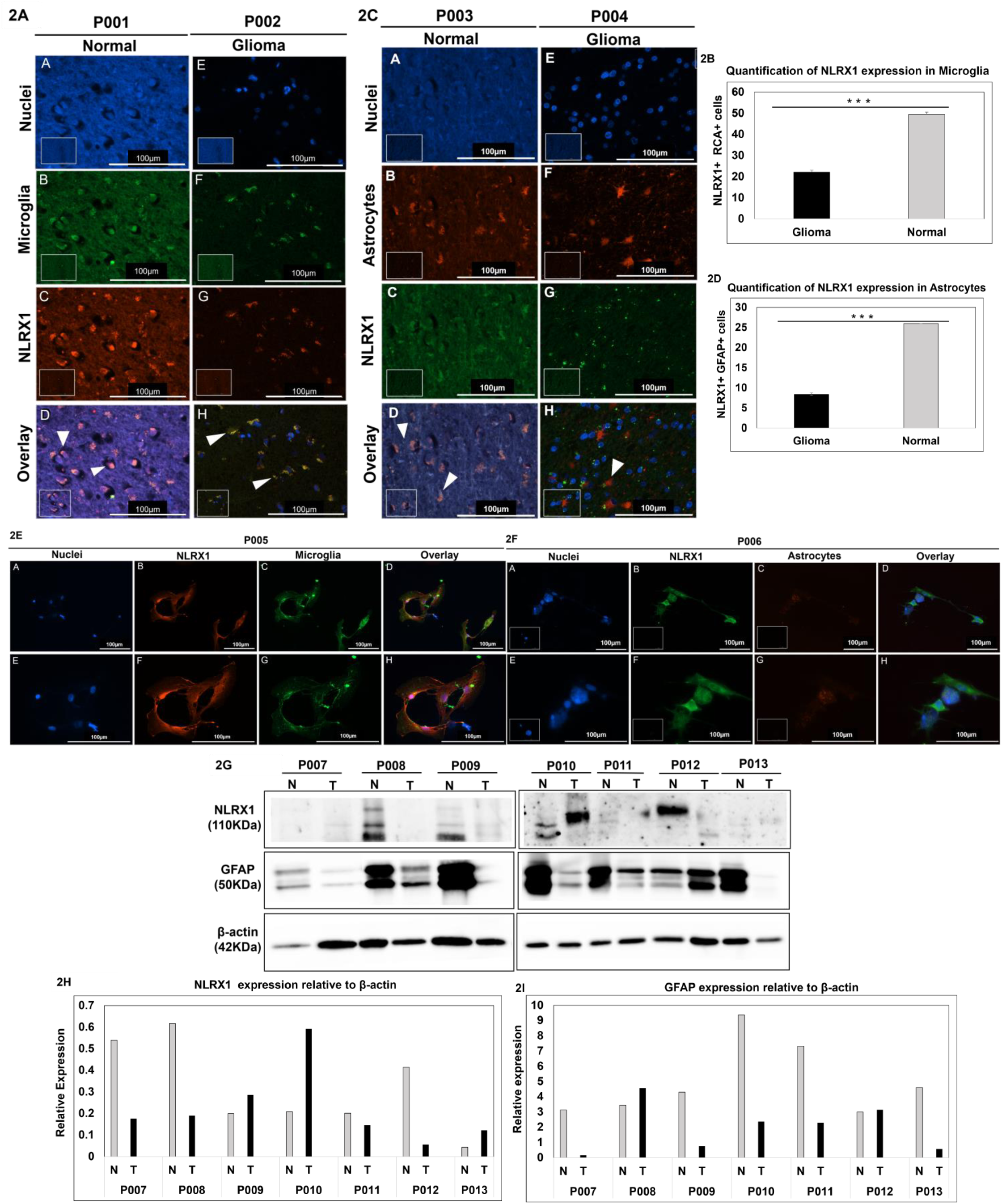
NLRX1 is differentially expressed across diverse glioma patients: A. Patient-derived paraffin-embedded glioma and normal brain tissue sections were stained with anti-NLRX1 antibody, and RCA lectin was used to detect microglia. The co-localization of NLRX1 and microglia can be visualized by the punctate red staining overlapping with green (arrowheads). DAPI was used to stain nuclei (blue). At least 7 frames were imaged per section. Images are representative of all frames. Inset represents primary antibody control and in overlays inset represents zoomed-in area. Scale bar, 100μm. B. The expression of NLRX1 is quantified across the microglia of normal and glioma tissue sections by manually blindly counting the NLRX1-expressing microglia. Data are represented as mean ± SEM. Error bars indicated SEM. ^∗^p < 0.05, **p < 0.01, ***p < 0.005 (Student’s t-test). C. Patient-derived paraffin-embedded glioma and normal brain tissue sections were stained with anti-NLRX1 antibody and GFAP was used to detect astrocytes. The co-localization of NLRX1 and astrocytes can be visualized by the punctate red staining overlapping with green (arrowheads). DAPI was used to stain nuclei (blue). At least 7 frames were imaged per section. Images are representative of all frames. Inset represents primary antibody control and in overlays inset represents zoomed-in area. Scale bar, 100μm. D. Expression of NLRX1 is quantified across Astrocytes of normal and glioma tissue sections by manual blind counting of NLRX1-expressing microglia. Data are represented as mean ± SEM. Error bars indicated SEM. ^∗^p < 0.05, **p < 0.01, ***p < 0.005 (Student’s t-test). E. Patient-derived tumor primary cells were stained for NLRX1 (red), RCA (green), and nuclei (DAPI). Inset represents primary antibody control, Scale bar, 100μm. At least 7 frames were imaged per well of the two well chamber slides. Images are representative of all frames. F. Patient-derived tumor primary cells were stained for NLRX1 (green), GFAP (red), and nuclei (DAPI). Inset represents primary antibody control, Scale bar, 100μm. At least 7 frames were imaged per well of the two well chamber slides. Images are representative of all frames. G. NLRX1 and GFAP expression was quantified across diverse glioma-grade patient tissues using western blot. β-actin is used as a loading control. 20μg protein sample is loaded in each well. N-Normal, T-tumor (n=7). H, I. The western blot data was quantified using densitometry.

**Table 1.** Details of patients.

| Patient number | Age/Sex | Pathology report | IDH status |
| --- | --- | --- | --- |
| P001 | - | Normal brain (Spinal Cord) | Unknown |
| P002 | - | Astrocytoma (WHO grade II glioma) | Unknown |
| P003 | - | Normal brain (Corpus Callosum) | Unknown |
| P004 | - | Astrocytoma (WHO grade II glioma) | Unknown |
| P005 | 39 Yr/M | Astrocytoma (CNS WHO Grade 2) | IDH-noncontributory |
| P006 | 34 Yr/F | Oligodendroglioma (CNS WHO Grade 2) | IDH1-noncontributory |
| P007 | 22/M | Central Neurocytoma, grade 2 (WHO 2016) | Unknown |
| P008 | 45/F | Astrocytoma, CNS WHO grade 3 (WHO CNS 2021) | IDH-mutant |
| P009 | 49/F | Neuroendocrine Carcinoma | Unknown |
| P010 | 48/M | Morphological features are of diffuse astrocytoma with gemistocytic and oligodendroglial areas, CNS WHO grade 2 (CNS WHO, 2021) | IDH-noncontributory |
| P011 | 32/M | Oligodendroglioma, WHO Grade 2 | IDH-noncontributory |
| P012 | 30/M | Diffuse Astrocytoma with oligodendroglial-like areas, CNS WHO grade 2 | IDH-noncontributory |
| P013 | 68/F | GBM, NOS, CNS classification 2021 | IDH-noncontributory |

### 3. *Nlrx1* knockdown attenuates GBM cell proliferation and migration

NLRX1 is a mitochondrial protein that regulates several cellular processes crucial for maintaining homeostasis. Recent studies have investigated the involvement of NLRX1 in maintaining metabolic homeostasis (Singh et al., 2019, 2015; Soares et al., 2014). In order to investigate the potential role of NLRX1 in the maintenance of GBM cells, we silenced the expression of *Nlrx1* in LN-229 cells using siRNA-mediated transfection (Fig EV1). Further, to check the potential influence of NLRX1 on cell proliferation, we performed a colony formation assay using LN-229 cells that were either wild-type (WT) or lacking the *Nlrx1* gene (Fig 3B, C, D). We observed a decrease in the rate of cell proliferation in *Nlrx1^-/-^* LN-229 cells. The *Nlrx1^-/-^* LN-229 cells exhibit a significantly lower colony count and smaller colony size compared to the WT cells (Fig 3C, D). Encouraged by these findings, we also investigated whether NLRX1 plays a direct role in the regulation of cell migration. In order to investigate the impact of NLRX1 on the migration of GBM cells, we conducted a wound healing assay utilizing WT and *Nlrx1^-/-^* LN-229 cells (Fig 3E, 3F) (Fig EV1). The scratch is imaged at 0 hours and 24 hours and the area of the scratch is quantified using ImageJ. Bright-field imaging and quantitative analysis of these images show that the area of scratch is greater in *Nlrx1^-/-^* LN-229 cells at 24 hours compared to WT cells. Thus, the migration rate varies between *Nlrx1^-/-^* LN-229 cells and WT cells, suggesting that NLRX1 potentially plays a role in regulating GBM cell, LN-229 migration.

**Figure 3.**
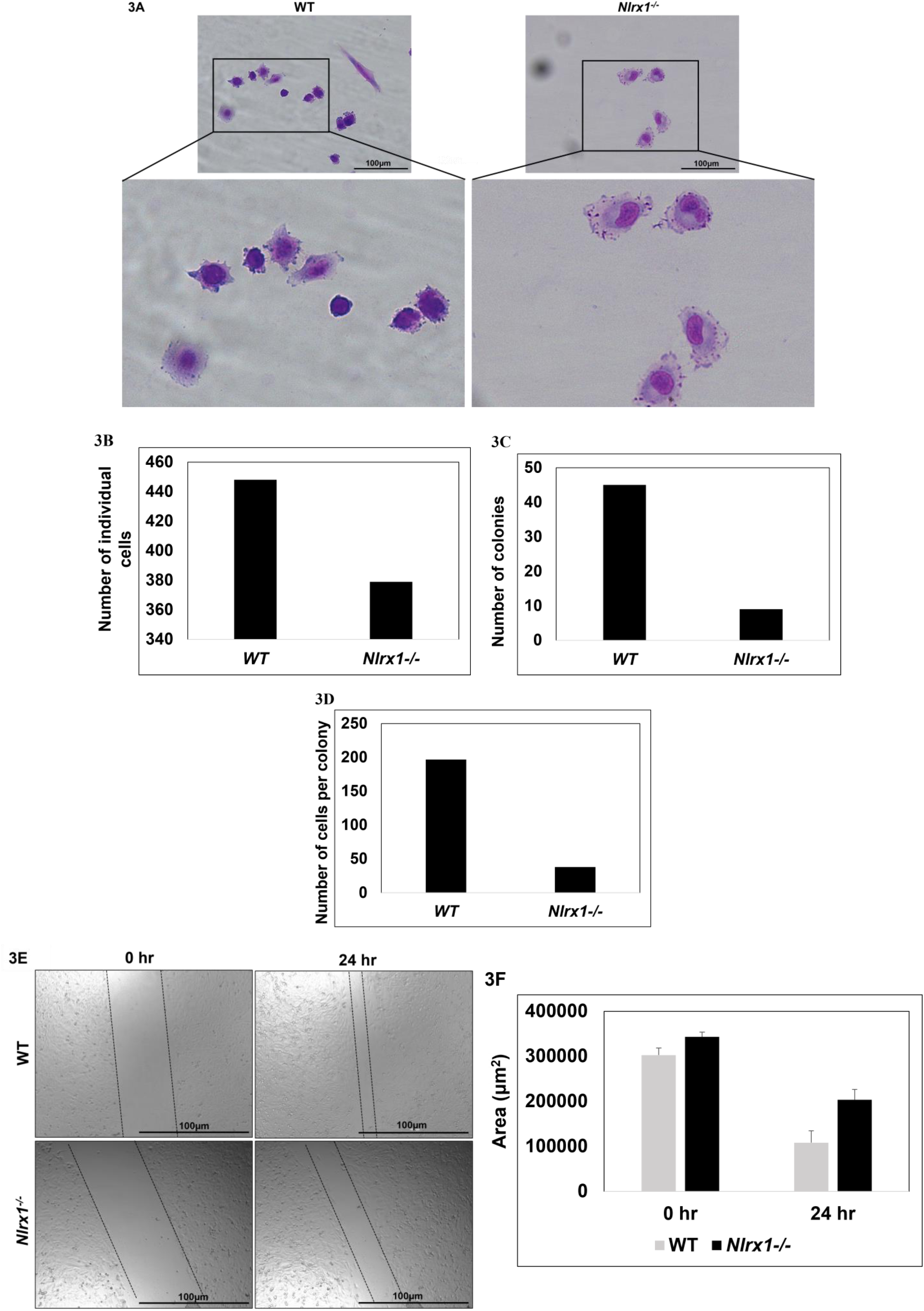
*Nlrx1* deficiency attenuates GBM cell proliferation and migration: A-B. colony formation assay was performed in WT and *Nlrx1^-/-^* LN-229 cells. A. WT and *Nlrx1^-/-^* LN-229 cells stained with Giemsa after 48 hours of seeding. At least 7 frames were imaged per well of the two well chamber slides, Scale bar, 100μm. Images are representative of 3 experiments. B, C, D. The number of individual cells, number of colonies, and number of cells per colony were counted in WT and *Nlrx1^-/-^* LN-229 cells by blind reader after 48 hours of cell seeding. Experiment was performed three times however; the graph presents data from one representative experiment. E, F. WT, and *Nlrx1^-/-^* LN-229 cells were imaged for 24 hours to assess the cell migration through wound healing assay. E. Images show the area of scratch at 0 hours and 24 hours in WT and *Nlrx1^-/-^* LN-229 cells. Scale bar, 100μm. Images are representative of 3 experiments. F. The area of scratch is quantified through Image J. The graph is representative of three experiments. Error bars indicated SEM.

### 4. TNT formation is increased in *Nlrx1* deficient GBM cells

An important phenomenon that cells use to exchange cytoplasmic factors and organelles is the formation of tubular connections, called tunneling nanotubes (TNTs) between cells. TNTs are cellular processes that enable cell-to-cell communication at distances from 30 to 500 µm (Ariazi et al., 2017). It has been seen that heterogeneous GBM cells express TNTs upon oxidative stress, and temozolomide (TMZ) and Ionizing Radiation (IR) treatment (Valdebenito et al., 2020). Further, it has been seen that glioma stem-like cells (GSLCs), grown in classical two-dimensional (2D) culture and three-dimensional (3D) tumor organoids, formed functional TNTs, which allowed mitochondria transfer (Pinto et al., 2021). Through TNTs, cells transport mitochondria and fulfill their energy requirements. siRNA-mediated silencing of *Nlrx1* in LN-229 cells was utilized to assess the contribution of NLRX1 to TNT formation since NLRX1 is an important regulator of cell metabolism (Fig EV1). These *Nlrx1^-/-^* LN-229 cells were cocultured with another GBM cell line, U87-MG, and assessed for TNT formation. As there is no known marker for TNTs, we performed electron microscopy to characterize TNT formation (Fig 4A). In addition, we conducted F-actin staining using phalloidin to stain TNTs formed between GBM cells. From the fluorescent images, we observed a higher number of TNTs in *Nlrx1^-/-^* GBM cells in comparison to WT cells (Fig 4B). In order to quantify the obtained observation, we imaged WT and *Nlrx1^-/-^* LN-229 cells co-cultured with U87-MG cells for 24 hours through continuous imaging. Fig 4C displays the representative images showing TNT formation at 0 and 24 hours in WT and *Nlrx1^-/-^* cells. It is evident that the number of TNTs is greater in *Nlrx1^-/-^* GBM cells in comparison to WT cells. The graph illustrates that at 0 hours, there were almost comparable numbers of TNTs. However, after 24 hours, there is a significant increase in the number of TNTs in *Nlrx1^-/-^* coculture (P<0.01) (Fig 4D). Thus, the loss of NLRX1 increases the number of TNTs between GBM cell lines, suggesting that NLRX1 regulates TNT formation. One possible explanation for these observations is that NLRX1-deficient LN-229 cells experience metabolic stress. Previous research has demonstrated that NLRX1 has a role in regulating the mitochondrial respiratory functions, ATP level, NAD+/NADH balance, and mitochondrial dynamics in triple-negative breast cancer cells in the presence of TNF-α (Singh et al., 2019). In the deficiency of NLRX1, an energy imbalance arises in LN-229 cells. To prevent this energy imbalance and maintain bioenergetic homeostasis, *Nlrx1^-/-^* LN-229 cells generate a higher number of TNTs with WT U87-MG cells, that have normal NLRX1 expression.

**Figure 4.**
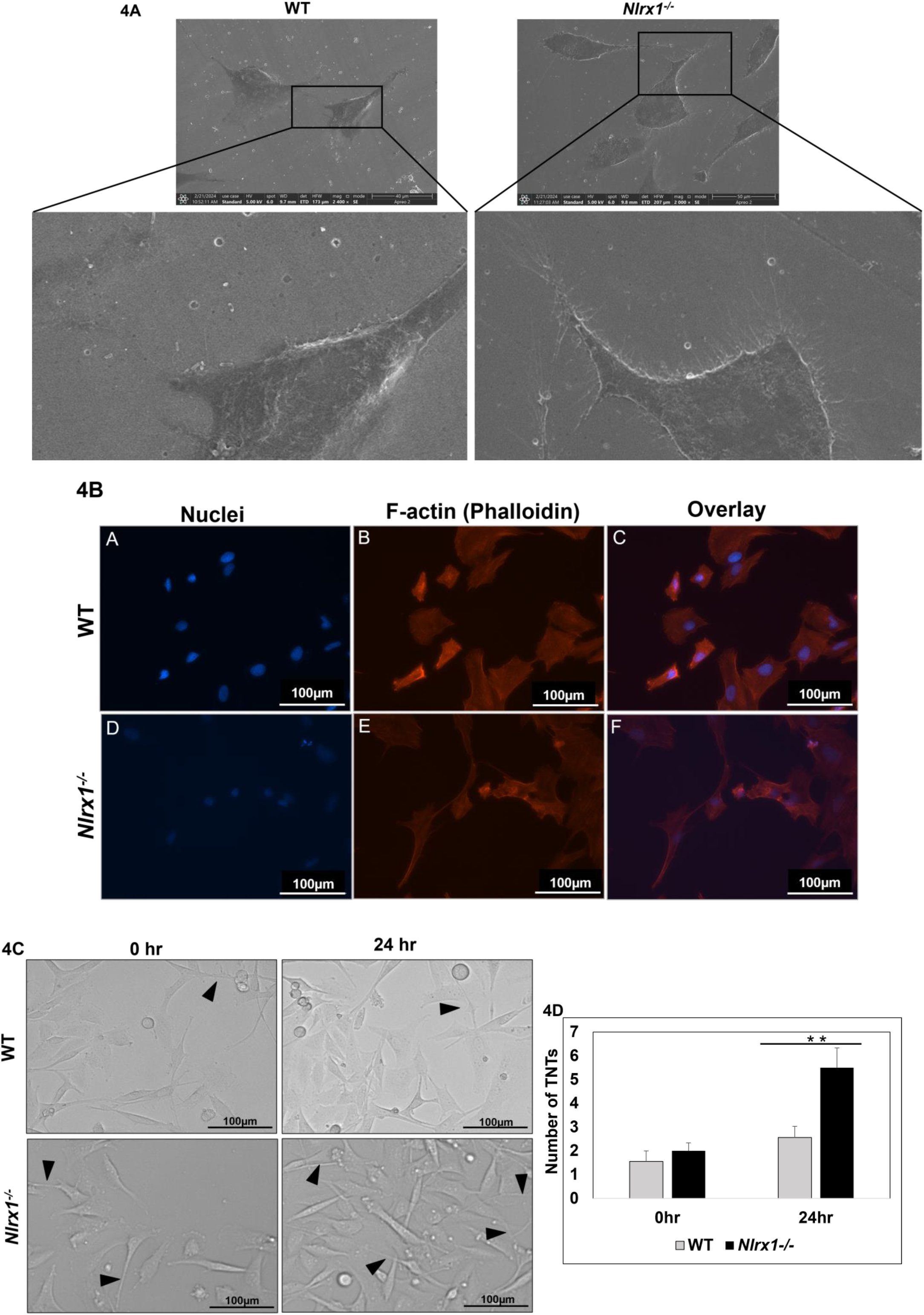
TNT formation is increased in *Nlrx1* deficient GBM cells: A. WT U87-MG cells were co-cultured with WT and *Nlrx1^-/-^* LN-229 cells and TNTs were analyzed using SEM. Images are representative of 2 experiments. B. LN-229 and U87-MG co-cultured cells were stained for F-actin (red), and nuclei (DAPI). At least 7 frames were imaged per well of the two well chamber slides. Images are representative of all frames. Inset represents Phalloidin control. Scale bar, 100 µm. C. LN-229 and U87-MG co-cultured cells were imaged for 24 hours to assess the TNT formation. Images are representative of 3 experiments. Scale bar, 100μm. D. The number of TNTs was quantified by manual blind counting of TNTs. The graph is representative of three experiments. Data are represented as mean ± SEM. Error bars indicate SEM. ^∗^p < 0.05, **p < 0.01, ***p < 0.005 (Student’s t-test).

### 5. 3D spheroid growth is attenuated in *Nlrx1* deficient GBM cells

Considering the role of NLRX1 in regulating cell proliferation and migration in 2D cultures we next investigated the role of NLRX1 on the 3D growth of tumors. Recent studies have shown that 3D cultures offer a physiologically relevant model over 2D cultures. Compared to conventional 2D culture, 3D tumor spheroids more closely resemble the GBM microenvironment *in vivo* in the context of spatial cell-cell and cell-extracellular matrix (ECM) interactions (Lin and Chang, 2008) (Mueller-Klieser, 1997). Spheroids are formed by the spontaneous aggregation of cells. After initial cell-cell contact, cells upregulate E-Cadherin, which accumulates on the cell surface, and then the spheroid becomes a compact structure through strong intercellular E-cadherin interactions (Lin et al., 2006). In order to investigate the potential involvement of NLRX1 in tumor formation, we utilized WT and *Nlrx1^-/-^* LN-229 cells to generate spheroids (Fig EV1). WT and *Nlrx1^-/-^* LN-229 cells were seeded and temporal live cell imaging was performed for 48 hours to create movies of the spheroid generation process (EV movie 1). Movies 1A and 1B show the spheroid formation of WT and *Nlrx1^-/-^* LN-229 cells, respectively. Furthermore, end-point day-wise imaging of spheroids was performed for the next five days (Fig EV2A). From the images, it can be noted that WT LN-229 cells formed larger spheroids compared to the *Nlrx1^-/-^* LN-229 cells (Fig 5A). To quantify this data, the area and diameter of spheroids from WT and *Nlrx1^-/-^* LN-229 were measured using ImageJ (Fig EV2B, C). Based on the graph (Fig 5B, C), it is evident that there is a significant difference in the size and diameter of WT and *Nlrx1^-/-^* LN-229 spheroids at day 5 (P<0.05) (P<0.005). The spheroids derived from *Nlrx1^-/-^* LN-229 cells exhibit reduced area and diameter. These discoveries are consistent with our findings in the 2D data, where deficiency of NLRX1 resulted in reduced cell proliferation. In addition, the circularity and compactness of WT and *Nlrx1^-/-^* spheroids were assessed to get insight into the impact of NLRX1 on tumor invasiveness (Fig 5D, E) (Fig EV2D, E). Compactness and circularity of a spheroid are used as a tool to measure the invasive potential of a spheroid (Lim et al., 2020). Recently, different image-based methods have been proposed for the quantitative analysis of cancer cell invasiveness. For example, Lim et al. developed an image analysis–based method to quantify the invasiveness of HT1080 human fibrosarcoma tumor cell spheroids (Lim et al., 2020). They segmented a cell-covered area into three subareas using objectively set threshold pixel intensities and calculated invasion indices using these subareas. Similarly, we also used an image-based approach to analyze the circularity and compactness of spheroids, as previously described (Leung et al., 2015). We noticed a difference in the circularity and compactness of WT and *Nlrx1^-/-^* spheroids, although the difference was not statistically significant (P>0.05). Thus, our data indicates that NLRX1 affects the proliferation of GBM cells in the 3D microenvironment, however, it does not affect the invasiveness potential of spheroids.

**Figure 5.**
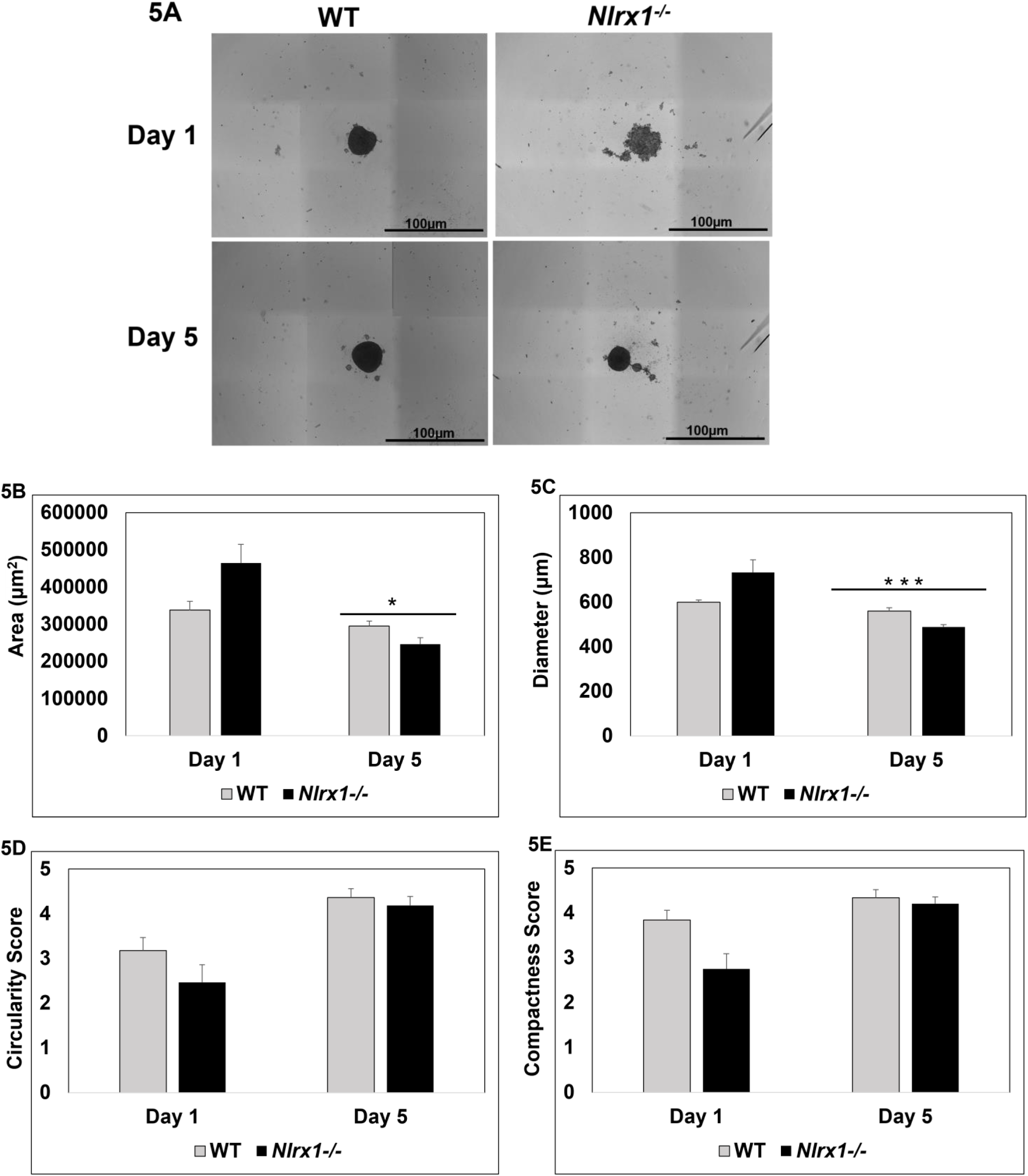
3D spheroid growth is attenuated in *Nlrx1* deficient GBM cells: WT and *Nlrx1^-/-^* LN-229 cells were seeded for spheroid formation and imaged for 7 days. A. Images of spheroids from WT and *Nlrx1^-/-^* LN-229 cells from day 1 and day 5. Images are representative of 3 experiments. Scale bar, 100μm. B-C. The area and diameter of spheroids from WT and *Nlrx1^-/-^* LN-229 cells were quantified using ImageJ. The graph is representative of three experiments. Data are represented as mean ± SEM. Error bars indicate SEM. ^∗^p < 0.05, **p < 0.01, ***p < 0.005 (Student’s t-test). D-E. The circularity and compactness of spheroids from WT and *Nlrx1^-/-^* LN-229 cells were analyzed using blind readers. The graph is representative of three experiments. Data are represented as mean ± SEM. Error bars indicated SEM.

### 6. NLRX1 regulates cell metabolism during energy crisis

Earlier studies have shown that NLRX1 acts as a metabolic regulator by regulating diverse metabolic processes, including glycolysis, oxidative phosphorylation, and mitochondria fission. Soares et al show a decreased expression of NLRX1 upon glucose starvation in mouse embryonic fibroblasts (MEFs) and linked this finding with NLRX1 being a regulator of glycolysis (Soares et al., 2014). Further, to elaborate on the role of NLRX1 in metabolic regulation, we treated GBM cells (LN-229) and astrocyte cells (SVG) with a glycolytic inhibitor 2-deoxyglucose (2-DG). NLRX1 expression analysis showed an increased NLRX1 expression in both cell types upon 2-DG treatment. However, the increase was significant in astrocytes compared to GBM cells (Fig 6A, B) (p=0.01). An important phenomenon that cells employ during energy crises or nutrient starvation to maintain metabolic homeostasis is autophagy (He et al., 2018). To confirm whether NLRX1 regulates autophagy in GBM, we utilized *Nlrx1^-/-^* and WT LN-229 cells and checked the expression of autophagy markers including LC3B (Microtubule-associated proteins 1A/1B light chain 3B), Beclin-1, and ATG5 (autophagy-related protein 5) (Fig 6C, D) (Fig EV1). The western blot shows a significant decrease in the expression of autophagy markers such as LC3BI (P=0.05), LC3BII (P<0.05), and Beclin-1 (P=0.05) in *NLRX1^-/-^* LN-229 cells. This data shows that NLRX1 is linked with autophagy maintenance and deficiency of NLRX1 leads to defects in the autophagy pathway (Fig 6C, D). Further, to check the expression of autophagy-related proteins in response to nutrient starvation, GBM, and astrocyte cells were subjected to glucose and amino acid starvation as previously described (Jusović et al., 2023; Karabiyik et al., 2021; Li et al., 2023; Martin et al., n.d.; Park et al., 2023). Depletion of glucose uptake leads to changes in protein expression from the autophagy pathway. However, the changes were not significant (P>0.05) (Fig EV4). This may be due to the fact that cells that were subjected to protracted glucose starvation developed resistance to these nutrient-depleting conditions and achieved metabolic homeostasis. In response to amino acid starvation, expression of ATG5 and NLRX1 increases until 8 hours; however, there is a consistent decrease in expression after 8 hours (Fig EV5). This data further confirms the interaction of NLRX1 with ATG5. This supports previous findings of NLRX1 interacting with ATG5 via mitochondrial Tu translation elongation factor (TUFM) in HEK293T cells (Lei et al., 2012a).

**Figure 6.**
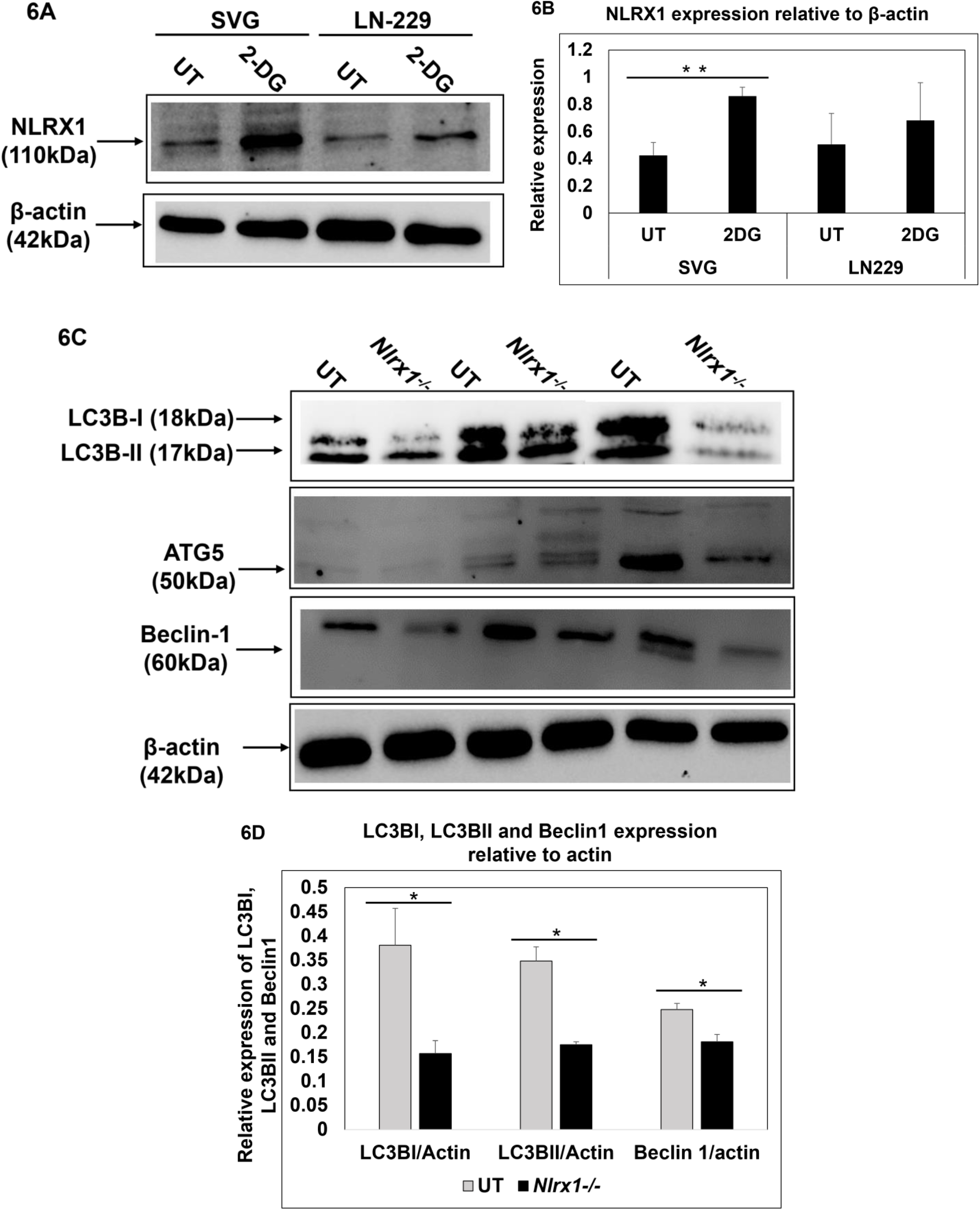
NLRX1 regulates cell metabolism during energy crisis: A. NLRX1 expression in Untreated, and 2-deoxy-glucose (2-DG) (10mM, 24hr) treated SVG and LN-229 cells was analyzed using western blot. β-actin is used as a loading control. 10µg protein sample is loaded in each well. Images are representative of 3 experiments. B. The western blot data was quantified using densitometry. The graph is representative of three experiments. Data are represented as mean ± SEM. Error bars indicated SEM. ^∗^p < 0.05, **p < 0.01, ***p < 0.005 (Student’s t-test). C. LC3B, Beclin1, and ATG5 expression in WT and *Nlrx1^-/-^* LN-229 cells were analyzed using western blot. β-actin is used as a loading control. 7µg protein sample is loaded in each well. Images are representative of 3 experiments. D. The western blot data was quantified using densitometry. The graph is representative of three experiments. Data are represented as mean ± SEM. Error bars indicated SEM. ^∗^p < 0.05, **p < 0.01, ***p < 0.005 (Student’s t-test).

## Discussion and conclusion

NLRX1 (nucleotide-binding domain and LRR-containing protein X, also known as CLR11.3, NOD5, or NOD9) was first discovered as a mitochondria-associated NLR that possesses a mitochondrial targeting sequence (MTS) encoded in the first 39 amino acids of the protein (Arnoult et al., 2009). NLRX1 consists of 975 amino acids and is ubiquitously expressed in mammalian tissues. NLRX1 may have cell and cancer context-dependent, tumor-promoting, or tumor-suppressing roles in different cancers (Nagai-Singer et al., 2023; Singh et al., 2019, 2015; Tattoli et al., 2016). NLRX1 regulates cellular signaling pathways such as the Nuclear factor-kappa B (NF-κB), Signal transducer and activator of transcription 3 (STAT3), Mitogen-activated protein kinase (MAPK), and Mitochondrial antiviral signaling protein (MAVS). In addition to these signaling pathways, NLRX1 also regulates several cellular processes, including inflammation, mitochondrial signaling, mitochondrial fragmentation, Reactive oxygen species (ROS) production, oxidative phosphorylation, autophagy, mitophagy, cell proliferation, and cell migration (Nagai-Singer et al., 2019). The expression pattern and role of NLRX1 have been studied in various cancers, including breast cancer, pancreatic cancer, and colorectal cancer (Nagai-Singer et al., 2023; Singh et al., 2019; Tattoli et al., 2016). However, the expression pattern of NLRX1 and how it regulates GBM has remained unexplored. Over the past decade, numerous significant findings have been related to the growth, metabolism, and signaling of gliomas. Nevertheless, the tumor-immune interactions in the tumor microenvironment remain largely unknown and necessitate additional investigation. Within malignant brain tumors, the cellular components of the microenvironment serve several roles, either promoting the growth of the tumor or preventing its malignant characteristics (Whiteside, 2008). Immune cells have been recognized as the predominant cell type in the cellular microenvironment. Over 95% of the cells from the myeloid population were identified as macrophages/microglia (Darmanis et al., 2017). Astrocytes regulate metabolic conditions and supply energy sources to neurons. Tumor-associated astrocytes and microglia play a crucial role in preserving the immunosuppressive TME and facilitating the invasion of tumor cells.

Given that NLRX1 functions as an innate immune receptor, we examined the expression pattern of NLRX1 in both innate immune cells and GBM cells. The subcellular localization of NLRX1 is cell- and microenvironment-dependent; however, it is predominantly present in mitochondria, cytoplasm, and cell membranes. We found differential expression of NLRX1 in GBM and normal cell types, as shown in Fig 1A. This may be because all these cell lines have different microenvironments associated with differential oncogenic signaling. In the GBM cell lines LN-229 and astrocyte cell line SVG, NLRX1 colocalizes with cytoskeletal components, actin, and mitochondria (Fig 1D, E). However, if we closely see the expression pattern of NLRX1 in LN-229 cells, it is highly expressed on cell membrane protrusions called lamellipodia. These structures are essential for cell migration and invasion. From this observation, we can assume that NLRX1 may have a role in regulating cell migration in LN-229 cells. SVGs lack this expression pattern, which may be due to their differential migration and invasion abilities as compared with the invasive GBM cell line, LN-229 cells. Endogenous NLRX1 expression was analyzed in astrocytes and microglia of glioma tissue sections. The number of NLRX1 expressing astrocytes and microglia is higher in normal brain tissue than in glioma tissues (P<0.005) (Fig 2A-2D). Further studies are required to see how the increased frequency of NLRX1-expressing microglia and astrocytes in normal tissues compared to glioma tissues is associated with the evolution of gliomas and immunosuppressive TME. Moreover, high NLRX1 expression was found in primary microglia and astrocytes isolated from fresh human glioma tissue samples (Fig 2E-2F). This data further indicates that NLRX1 has a role in regulating innate immunity in glioma and, ultimately, tumor progression. Gliomas are highly heterogeneous due to differences in the oncogenic signaling in different cell populations (Bianco et al., 2017). Interestingly, we found that NLRX1 expression also varies from patient to patient in the same grade of glioma and across different grades (fig 2G, 2H). We also observed a differential NLRX1 expression pattern between normal and tumor tissues. This data shows the differential NLRX1 signaling in the normal brain versus gliomas. Earlier studies have shown that NLRX1 regulates tumor progression by controlling the ability of cancer cells to proliferate and migrate. NLRX1 can positively or negatively affect the proliferation and migration of cancer cells depending on the cell type and microenvironment (Nagai-Singer et al., 2023; Singh et al., 2019, 2015). We have shown that the loss of NLRX1 decreases the proliferation and migration of GBM cell line, LN-229 cells (Fig 3). The data we obtained aligned with the findings of a previous study on invasive breast cancer cell lines (Singh et al., 2019).

Additionally, it was noted that the quantity of TNTs is more significant in cells lacking NLRX1 (*Nlrx1^-/-^)* compared to WT cells (P<0.01) (Fig 4). TNTs are tubular connections between cells that help them maintain metabolic homeostasis by enabling the exchange of subcellular organelles such as mitochondria and endoplasmic reticulum (Venkatesh and Lou, 2019). In addition, we saw a decline in cell proliferation in the 3D microenvironment, as the spheroids created from *Nlrx1^-/-^* LN-229 cells exhibited smaller area and diameter compared to spheroids from normal cells (P<0.05) (P<0.005) (Fig 5A-C). Therefore, our two-dimensional cell culture data and three-dimensional cell culture data both show reduced proliferation in *Nlrx1^-/-^* LN-229 cells. In addition, we assessed the circularity and compactness of the tumor cell spheroids to evaluate the invasive potential of GBM cell lines. Our observation revealed no significant difference in the circularity and compactness of spheroids from *Nlrx1^-/-^* LN-229 cells and WT cells (P>0.05) (Fig 5D, E). As these are subjective image-based scoring assessments, an in-depth cellular and molecular assessment, beyond the scope of this manuscript, will be needed to assess the role of NLRX1 in the invasiveness of GBM cell lines.

Earlier studies have shown that NLRX1 expression is associated with glycolysis, as inhibition of glycolysis decreases NLRX1 expression in primary MEFs (Soares et al., 2014). Tumors have enhanced glycolysis to fulfill increased energy demands of cancer cells, also known as the ‘Warburg effect’ (Lebelo et al., 2019). An essential function of NLRX1 is to regulate cell metabolism. One study found that when NLRX1 was suppressed in HeLa cells, it reversed the acidification of the culture medium (Singh et al., 2015). This suggests that NLRX1 may play a role in the metabolic shift to glycolysis in tumor cells. NLRX1 also controls the functions of mitochondrial complexes I and III to sustain ATP levels in the presence of TNF-α and facilitate the metabolic shift towards aerobic glycolysis (Singh et al., 2019). To understand the role of NLRX1 in the metabolism of astrocytes and GBM cells, cells were treated with the glycolytic inhibitor 2-DG, and the expression of NLRX1 was analyzed (Fig 6A, B). Increased expression of NLRX1 upon 2-DG treatment further confirms the role of NLRX1 in maintaining metabolic homeostasis. As NLRX1 helps in metabolic homeostasis maintenance, increasing NLRX1 expression by cells is an attempt to gain metabolic equilibrium. However, the GBM cells are much more robust in comparison to astrocytes because they have altered oncogenic signaling pathways, i.e., LN-229 cells have p53 gene mutation (“BPA-9-469,” n.d.; Schlapbach and Fontana, 1997), which leads to increased amino acid metabolism, and lipid synthesis (Laue et al., n.d.; Tombari et al., 2023). Additionally, in conditions of glucose starvation, GBM cells can fulfill their energy demands by the Wnt/β-catenin signaling and glutaminolysis pathway (Stuart et al., 2023; Yusuf et al., 2022). Thus, these altered signaling in GBM cells helps them to survive and sustain their metabolism in adverse conditions. Contrary to GBM cells, astrocytes do not have any modified signaling. They are crucial in sustaining neuronal metabolism by supplying pyruvate or lactate to neurons for energy maintenance (Almeida et al., 2023; Lee et al., 2016). Therefore, astrocytes exhibit greater susceptibility to a disturbance in metabolic homeostasis caused by glycolytic inhibitors than GBM cell lines, leading to a more pronounced rise in NLRX1 expression in astrocytes (P=0.01). Previous research has demonstrated the context and environment-dependent interaction between NLRX1 and diverse autophagy-related proteins. NLRX1 regulates autophagy by interacting with the ATG5-ATG12 (autophagy-related5-autophagy-related12) complex through the TUFM domain, the LC3 protein through the LRR domain of NLRX1, and the Beclin1-UVRAG complex (Aikawa et al., 2018; Lei et al., 2012b; Zhang et al., 2019). Autophagy may have both tumor-promoting and tumor-suppressing effects in GBM (Meena and Jha, 2024). To test the hypothesis that NLRX1 affects autophagy in GBM, we utilized *Nlrx1*^-/-^ and WT LN-229 cells and assessed the expression of autophagy markers (Fig 6C, D). This analysis showed defective autophagy in *Nlrx1*^-/-^ LN-229 cells, proving that NLRX1 has a role in the regulation of autophagy in GBM cells.

Our study identifies the role of NLRX1 in the modulation of GBM pathophysiology. NLRX1 is one of the cellular metabolic regulators in GBM as it influences TNT formation, autophagy, and its expression changes in adverse metabolic conditions. Therefore, the cells have experienced a disruption in metabolic equilibrium under NLRX1-deficient conditions, resulting in decreased cell proliferation and migration. In order to maintain metabolic equilibrium or meet their energy demands, cells may have augmented the quantity of TNTs. Thus, cells may have increased the number of TNTs in order to survive the disrupted metabolic status. Gaining a more comprehensive understanding of this mechanism would facilitate the specific targeting of NLRX1 in order to govern the survival and metabolism of GBM cells. Multiple clinical trials are now investigating NLR-related compounds in different diseases (Wang et al., 2024). For example, specific NLRP3 inhibitors, such as DFV890, OLT1177, IFM Tre, Inzomelid, and Somalix are in different phases of clinical trials. DFV890 has completed a Phase I trial for coronavirus disease 2019 (COVID-19) pneumonia, OLT1177 has concluded a Phase IB trial, in which it improved left ventricular ejection fraction in patients with heart failure and reduced ejection fraction. Additionally, IFM Tre, Inzomelid, and Somalix have completed phase I clinical trials, while some other NLRP3 inhibitors are still under phase I trials, such as NT-0167, NT-0796, and ZYIL1. Similarly, NLRX1 agonist NX-13 has completed phase IB clinical trials (Verstockt et al., 2024). NX-13 activation of NLRX1 reduces intracellular reactive oxygen species and decreases inflammation in animal models of colitis. A phase 1A trial demonstrated a gut-selective pharmacokinetic profile with good tolerability. This phase IB trial evaluated the safety, tolerability, and pharmacokinetics of NX-13 in patients with active ulcerative colitis [UC]. There are a variety of potential outcomes associated with molecules that regulate the expression of the NLR. We believe our understanding of NLRX1 in GBM pathophysiology paves the possibility for the development of future GBM-targeting therapeutics that may delay disease progression and/or improve survival.

## Material and Methods

### 1. Cell culture

The human glioblastoma cell line LN-18 was purchased from the American Type Culture Collection (ATCC), and human glioblastoma cell lines LN-229 and U87-MG were purchased from the Cell Repository of the National Centre for Cell Science (NCCS) Pune, India. Human astrocyte cell line SVG was kindly shared by Dr Pankaj Seth from the National Brain Research Centre (NBRC), India. Cells were grown in media containing DMEM (HiMedia, AL151A-500ML) along with 10% FBS (Cell Clone, CCS-500-SA-U) and 1% antibiotic-antimycotic solution (penicillin, streptomycin, amphotericin B) (HiMedia, A002-200ML) at 37°C with 5% CO_2_.

### 2. Immunocytochemistry

Cells were seeded at 50,000 cells per well in cell culture medium in chamber slides and incubated in a CO_2_ incubator (5% CO_2_, 37°C temperature, and 95% humidity). Cells were washed with PBS, fixed using 4% paraformaldehyde (HiMedia, MB059-500ML) for 10 minutes, permeabilized with 0.1% triton-x (Sigma, 1001723790) in PBS (permeabilization buffer) for 15 minutes, and blocked with blocking buffer containing 5% FBS in permeabilization buffer for 1hr at 4°C (in a humidified chamber). Cells were immunolabelled with primary antibody (NLRX1: Novus biologicals, rabbit antibody, NBP1-76287, actin: Santa-Cruz Biotechnology, mouse antibody, sc-47778, GFAP: Cell signaling technology, mouse antibody, 3670S) and incubated overnight at 4°C. Next day, cells were washed with PBS, and incubated with secondary antibody (Alexa fluor 488nm: Invitrogen, rabbit antibody, A11008, Alexa fluor 594nm: Invitrogen, mouse antibody, A11005) for 1hr at 37°C (humidified and dark conditions), and slides were mounted (Sharma et al., 2019).

### 3. Immunohistochemistry

Immunohistochemistry was performed on 5-μm paraffin-embedded sections that were deparaffinized and rehydrated through alcohols as described previously (Sharma et al., 2019). The paraffin-embedded paraformaldehyde fixed glioma and normal brain tissue were obtained with approval from the Internal Review Board and the Ethics Committees of All India Institute of Medical Sciences (AIIMS), Jodhpur, and Tata Memorial Cancer Hospitals. Informed consent was acquired from human participants for the use of tissue samples for experiments. We have performed all experiments in accordance with the ethical guidelines and regulations of All India Institute of Medical Sciences (AIIMS), Jodhpur, and Indian Institute of Technology Jodhpur. To detect astrocytes, deparaffinized and rehydrated sections were incubated with 0.1% Triton/PBS for 20 min at RT, followed by 5% FBS in 0.1% Triton/PBS for 1 hour at 4°C. Subsequently, the sections were washed and incubated with primary antibody (NLRX1: Novus biologicals, rabbit antibody, NBP1-76287, Glial fibrillary acidic protein (GFAP) for detection of astrocytes: Cell signaling technology, mouse antibody, 3670S) overnight at 4°C in a humidified chamber. After washing with PBS, sections were incubated with a secondary (Alexa fluor 488nm: Invitrogen, rabbit antibody, A11008, Alexa fluor 594nm: Invitrogen, mouse antibody, A11005) for 1hr at 37°C (in a humidified, dark chamber). Subsequently, tissue sections were washed and mounted as previously described (Agrawal et al., 2021). To detect microglia, deparaffinized and rehydrated, sections were incubated with 0.1% Triton/PBS for 20 min at room temperature (RT). Subsequently, the sections were washed and incubated with RCA (Ricinus communis agglutinin) for detection of microglia (Vector laboratories, FL-1081) for 1 hour at 37°C in a humidified chamber. After washing with PBS, tissue sections were incubated in 5% FBS in 0.1% Triton/PBS for 1 hour at 4°C. Subsequently, the sections were washed and incubated with anti-NLRX1 primary antibody overnight at 4°C in a humidified chamber. After washing with PBS, sections were incubated with a secondary antibody (Alexa fluor 594: Life Technologies, rabbit antibody, A11012) for 1hr at 37°C (in a humidified, dark chamber). Subsequently, tissue sections were washed and mounted as previously described (Agrawal et al., 2021).

### 5. MitoTracker Staining

Human glioblastoma cell line, LN-229 cells, and human astrocyte cell line, SVG cells were seeded in chamber slides with growth medium and incubated in a CO_2_ incubator (5% CO_2_, 37°C temperature, and 95% humidity). At 60-80% confluency, media was removed from wells, and serum-free DMEM was added. Mito Tracker selectively stains live mitochondria in cells. Mito Tracker (12.5nM, ThermoFisher, 89874) staining was carried out as per manufacturer protocol. Cells were washed and fixed with paraformaldehyde (HiMedia, MB059-500ML), followed by permeabilization and blocking as described previously. Cells were immunolabelled overnight with anti-NLRX1 primary antibody. Cells were washed with PBS, incubated with a secondary antibody (Alexa fluor 594: Life Technologies, rabbit antibody, A11012), and slides mounted using FluoroshieldTM with DAPI (4’,6-diamidino-2-phenylindole) (Sigma, F6057-20ML) stained nuclei blue, as described previously (Tattoli et al., 2008(8).

### 6. Isolation of primary cells from patient tissues

The glioma tissues were obtained with approval from the Internal Review Board and the Ethics Committees of AIIMS, Jodhpur. Informed consent was acquired from human participants for the use of tissue samples for experiments. We have performed all experiments in accordance with the ethical guidelines and regulations of AIIMS, Jodhpur, and Indian Institute of Technology Jodhpur. Tissue sections were collected in aCSF (2mM CaCl_2_.2H_2_O, 10mM glucose, 3mM KCl, 26mM NaHCO_3_, 2.5mM NaH_2_PO_4_, 1mM MgCl_2_.6H_2_O, 202mM sucrose) at AIIMS Jodhpur. Once received at our laboratory, aCSF was discarded, and the tissue was weighed. Tissues were washed in aCSF and PBS. Then, tissues were chopped into approximately 1mm size pieces and transferred into a solution of 0.25% trypsin with EDTA. The tissue was dissociated by shaking at 250 RPM for 30 minutes at 37°C. After dissociation, neutralizing media was added to neutralize the trypsin, and the sample was centrifuged. The cell pellet is collected and resuspended in 1ml of culture medium (45% DMEM, 45% F12-nutrient medium, and 10% FBS). Cells were plated in a tissue culture flask and incubated in a CO_2_ incubator (5% CO_2_, 37°C temperature, and 95% humidity) (Agrawal et al., 2020).

### 7. Isolation of Mitochondria from cells

Mitochondria and cytosolic fractions were separated from the human GBM cell line, LN-229 cells, and human astrocyte cell line, SVG cells using the manufacturer’s protocol (Mitochondrial isolation kit, Thermofisher, 89874). Isolated mitochondria were stored in RIPA buffer for further experiments.

### 8. Western blotting

Cells or tissues were lysed in radioimmunoprecipitation assay (RIPA) buffer with freshly added protease inhibitors for 4 min at 4°C. The protein concentration was assessed using a Bradford assay. 10µg of protein was loaded onto SDS-PAGE and then transferred onto a nitrocellulose membrane, which was incubated with blocking buffer (5% skimmed milk) for 1.5 hours, followed by primary antibodies (NLRX1: Novus biologicals, rabbit antibody, rabbit antibody, NBP1-76287, TOM20: Santa-Cruz Biotechnology, mouse antibody, sc-17764, LC3B, ATG5, Beclin1: Novus biologicals, rabbit antibody, NB910-94877, ATG9: β-actin: Santa-Cruz Biotechnology, mouse antibody, sc-47778) overnight and appropriate secondary HRP-conjugated antibodies (Anti-rabbit IgG HRP-linked: Cell signaling technology, 7074S, Anti-mouse IgG HRP linked: Cell signaling technology, 7076S). Protein expression was visualized using the Azure Biosystems Gel Documentation system. ImageJ software was used for the densitometry analysis of the blots. The relative protein quantity was normalized to β-actin (Xu et al., 2020).

### 9. siRNA-mediated gene silencing

LN-229 cells were seeded in 6-well plates. After cell adherence, cells were left untreated (ctrl) or transfected with non-specific scrambled RNA control (scRNA 50nM) or *Nlrx1*-specific siRNAs (50 and 100 nM) in Opti-MEM medium. After 48 hours of treatment, the protein was isolated from transfected and control cells using RIPA buffer, and a western blot was performed to check gene silencing (Sharma et al., 2019).

### 10. Wound healing assay

WT and *Nlrx1*^-/-^ LN-229 cells were seeded in a 96-well plate (10,000 cells/well). After almost 90% confluency, a scratch was made in the well, and cells were observed for the next 24 hours. Images were taken using a multimode reader (cytation5, Agilent).

### 11. Tunneling nanotubes (TNT) visualization and quantification

For TNT analysis, *Nlrx1*^-/-^ LN-229 cells were co-cultured with U87-MG (1:1 ratio) in 96-well plates (2500 cells/well). Confluence used was 50 to 70 % to enable TNT extension and communication and reduce the possibility of overgrowth that can compromise TNT identification and characterization. The cells were imaged in a multimode reader (cytation5, Agilent) at 37°C temperature and 5% CO_2_ to avoid any significant variations in temperature or CO_2_. Cells were imaged for 24 h, recording every 30 min (Valdebenito et al., 2020).

### 12. Colony formation assay

WT and *Nlrx1*^-/-^ LN-229 cells were seeded at a density of 2500 cells per well (5% CO_2_; 37°C) in a chamber slide, and small colonies were observed after 36–48 hours. Colonies were stained with Giemsa, as previously described (Sharma et al., 2019). Slides were observed using a bright field microscope, and results were quantified by counting the number of colonies formed per well and cells present per colony.

### 13. Spheroid formation

WT and *Nlrx1*^-/-^ LN-229 cells were seeded in low attachment U-shaped 96 well plate for spheroid generation. Cells were maintained in a medium containing 0.1% antibiotic,10% FBS, and nutrient media (DMEM) at 37°C with 5% CO_2_. Cells began to aggregate within a few hours and formed spheroids within one day. The spheroids were imaged with a multimode reader (cytation5, Agilent).

### 14. Image acquisition of spheroid

The imaging of seeded spheroids was performed through an automated multimode reader (cytation5, Agilent) from day zero at 5% CO_2_ concentration and 37°C. Images were taken at different Z-planes, and after the imaging was completed, the Z-plane images were stitched using the built-in software. Final images were saved for further analysis.

### 15. Quantification of area and diameter of spheroids

The NIH Image J tool was used to quantify the spheroid diameter and area. The area of the spheroid was manually selected, and the area of the selected area was quantified using the Image J application. The diameter of the spheroid was measured along 2 perpendicular axes – horizontal and vertical. The diameter of a spheroid is taken as the average diameter of the horizontal and vertical axis of the spheroid (Leung et al., 2015).

### 16. Quantification of circularity and compactness of spheroids

To assess the circularity and compactness of the spheroid, blinded observers were asked to score each category on a 5-point scale. In each category, the lack of a visible spheroid or formation of multiple spheroids was assigned a score of 1. For compactness, spheroid with clearly visible spaces and gaps were classified as a loose aggregate (score = 2), and aggregates with no gaps within the cell mass but presenting diffuse borders were scored as a tight aggregate (score = 3). Further compaction led to the formation of distinct dark borders around spheroids with few loose cells attached; these were classified as compact spheroids (score = 4). At the most compact stage, cells on the surface of the spheroid were remodelled and followed the contour of the spheroid, creating a smooth and defined outline, and were classified as tight spheroids (score = 5). For circularity, spheroids with similar degrees of concave and convex outline were classified as irregular (score = 2). In contrast, those mainly consisting of convex borders but with small concave dimples were classified as minor irregular (score =3). Spheroids that are elongated with no concave outline sections receive a score of 4, and finally, symmetrically circular spheroids receive a score of 5 (Leung et al., 2015).

### 17. Scanning electron microscopy (SEM)

Cells were seeded on L-lysine-coated coverslips. When the cells adhered to the coverslip, they were fixed using 2.5% glutaraldehyde for 2 hours at 4°C. After fixation, cells were washed with 1X-phosphate buffer and then dehydrated with a graded ethanol series by subsequent exchanges of the following dilutions in distilled water – 50%, 70%, 95%, and 100% ethanol (15 minutes). After dehydration, cells were dried, coated with gold particles, and imaged using SEM (Fischer et al., 2012).

### 18. Phalloidin Staining

WT and *Nlrx1*^-/-^ LN-229 cells were seeded in a chamber slide with growth medium and incubated in a CO_2_ incubator (5% CO_2_, 37°C temperature, and 95% humidity). Cells were washed with PBS, fixed using 4% paraformaldehyde (HiMedia, MB059-500ML) for 10 minutes, and permeabilized with 0.1% triton-x (Sigma, 1001723790) in PBS for 15 minutes. Cells were stained with Rhodamine phalloidin (Invitrogen, R415) solution in 1% BSA solution (1:200) for 1 hour at RT. Cells were washed with PBS and mounted using Fluoroshield^TM^ with DAPI (4’,6-diamidino-2-phenylindole) (Sigma, F6057-20ML) (Pinto et al., 2021).

### 19. Statistical Analyses

Data are expressed as mean ± SEM. Unpaired Student’s t-tests were used to statistically evaluate significant differences. Differences were considered statistically significant if p<0.05. The student’s t-test was performed to determine significant group differences.

## Author Contribution

D.M. designed and performed the experiments (data analysis, immunofluorescence, western blotting, mitochondria isolation, SEM, primary cell isolation) and prepared and edited the initial manuscript draft. D.S. quantified the TNTs and calculated the area and diameter of spheroids. S.R. helped with cell culture and protein isolation-related experiments. N.B. quantified the spheroid’s circularity and compactness and quantified the IHC data. P.S. and S.C. helped with CFA quantification. V.J., M.G., and J.S.G. provided patient tissue samples for IHC, ICC, and western blot. S.J. conceptualized the study, designed experiments, and edited and reviewed the manuscript. All authors reviewed the manuscript.

## Acknowledgement

S.J.’s laboratory was established with institutional grants from the Indian Institute of Technology Jodhpur (IITJ) and is funded by the Ministry of Electronics and Information Technology, Government of India (No.4 (16)/2019-ITEA. Dr Pankaj Seth from the National Brain Research Centre (NBRC) kindly shared the human astrocyte cell line SVG. Human brain tissue sections were obtained from the All-India Institute of Medical Sciences (AIIMS) Jodhpur and Tata Memorial Hospital, Tata Memorial Center, Mumbai, Maharashtra, India. We thank Mr. Bharat Pareek, Technical Superintendent, Indian Institute of Technology Jodhpur, for his technical support in our lab.

## Disclosure and competing interest statement

The authors declare that they have no conflict of interest

## Funding Information

The work is funded by grants from the Ministry of Electronics and Information Technology, Government of India (No. 4 (16)/2019-ITEA). D.M. is supported by a fellowship from the Council of Scientific & Industrial Research (CSIR)-NET-JRF (09/1125(0018)/2021-EMR-I).

## Expanded View Figure legend

**Fig EV1.**
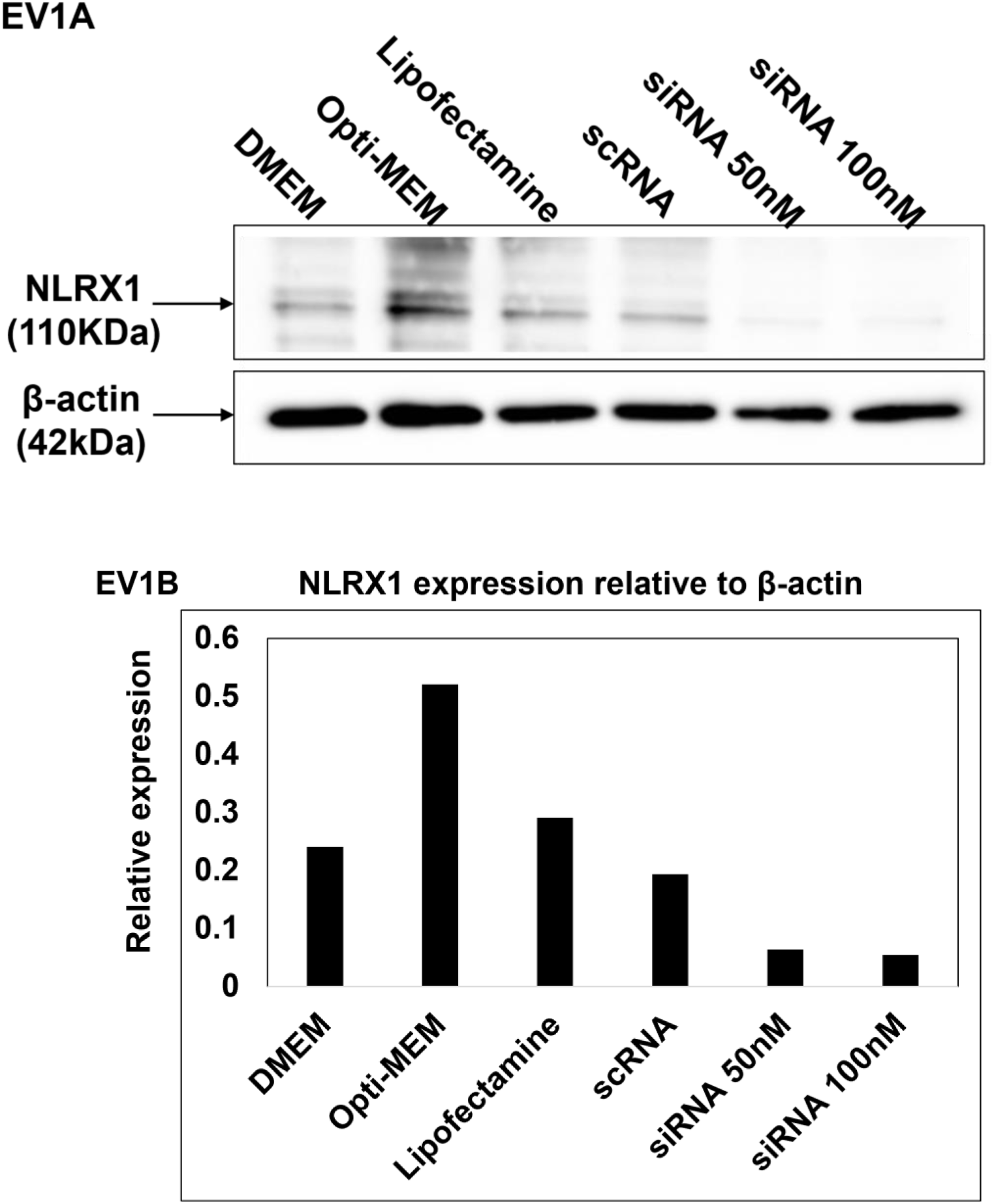
Knockdown of *Nlrx1* in GBM cell line, LN-229 using siRNA, related to Figures 3, 4, 5, and 6: A. NLRX1 expression was quantified in *Nlrx1* siRNA-treated LN-229 cells using a western blot. β-actin is used as a loading control. 15µg protein sample is loaded in each well. Images are representative of 3 experiments. B. The western blot data was quantified using densitometry. The graph represents data from one representative experiment.

**Fig EV2.**
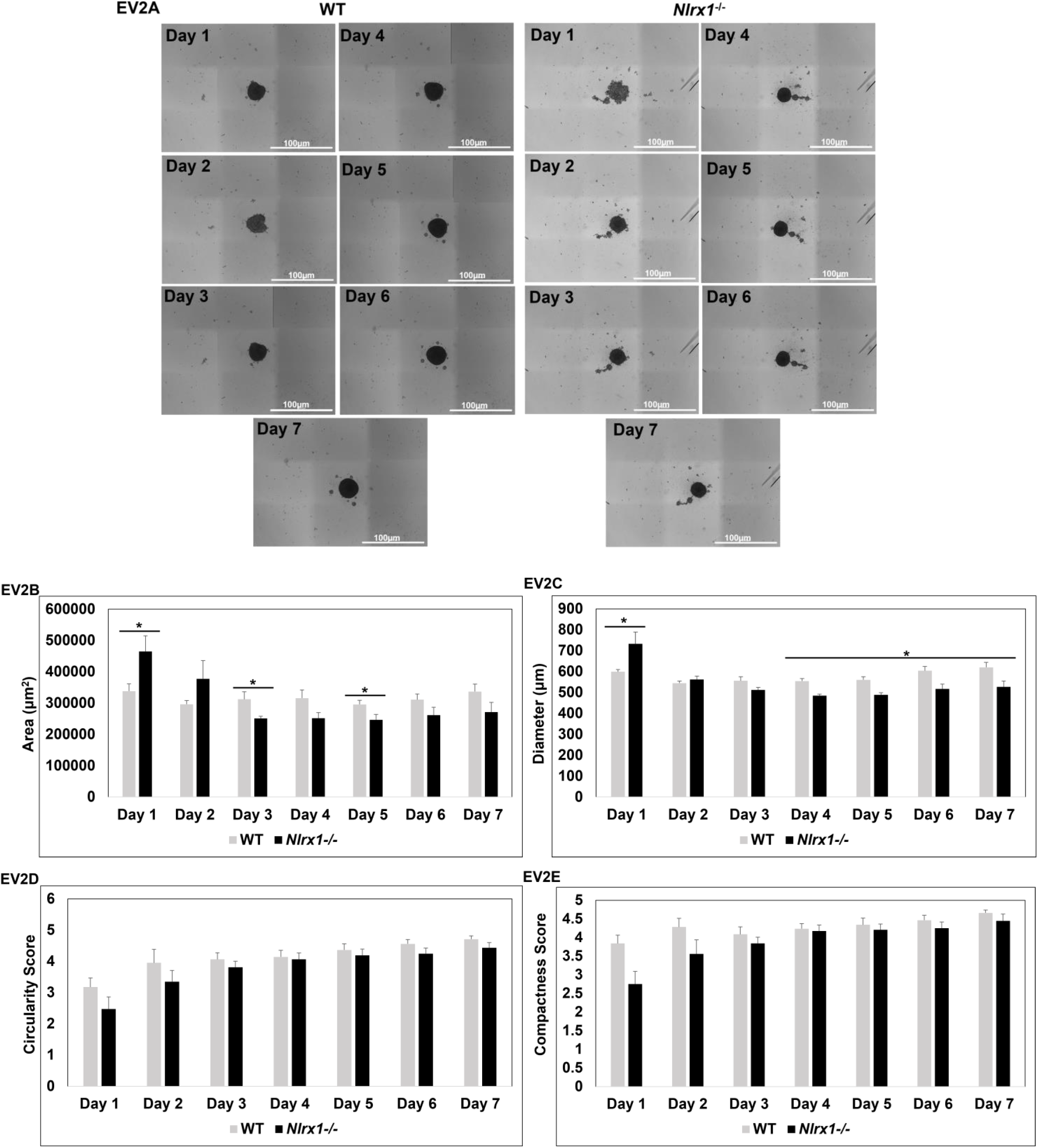
*Nlrx1* knockdown attenuates the capability of GBM cell lines to form spheroids in 3D microenvironment, related to Figure 5: WT and NLRX1^-/-^ LN-229 cells were seeded for spheroid formation and imaged for 7 days. A. Images of spheroids from WT and NLRX1^-/-^ LN-229 cells from day 1 to day 7. Images are representative of 3 experiments. Images were taken at a 4X objective lens, Scale bar, 100μm. B-C. Area and diameter of spheroids from WT and NLRX1^-/-^ LN-229 cells were quantified using ImageJ. The graph is representative of three experiments. Data are represented as mean ± SEM. Error bars indicated SEM. ^∗^p < 0.05, **p < 0.01, ***p < 0.005 (Student’s t-test). D-E. Circularity and compactness of spheroids from WT and NLRX1^-/-^ LN-229 cells were analyzed using blind readers. The graph is representative of three experiments. Data are represented as mean ± SEM. Error bars indicated SEM.

**Fig EV3.**
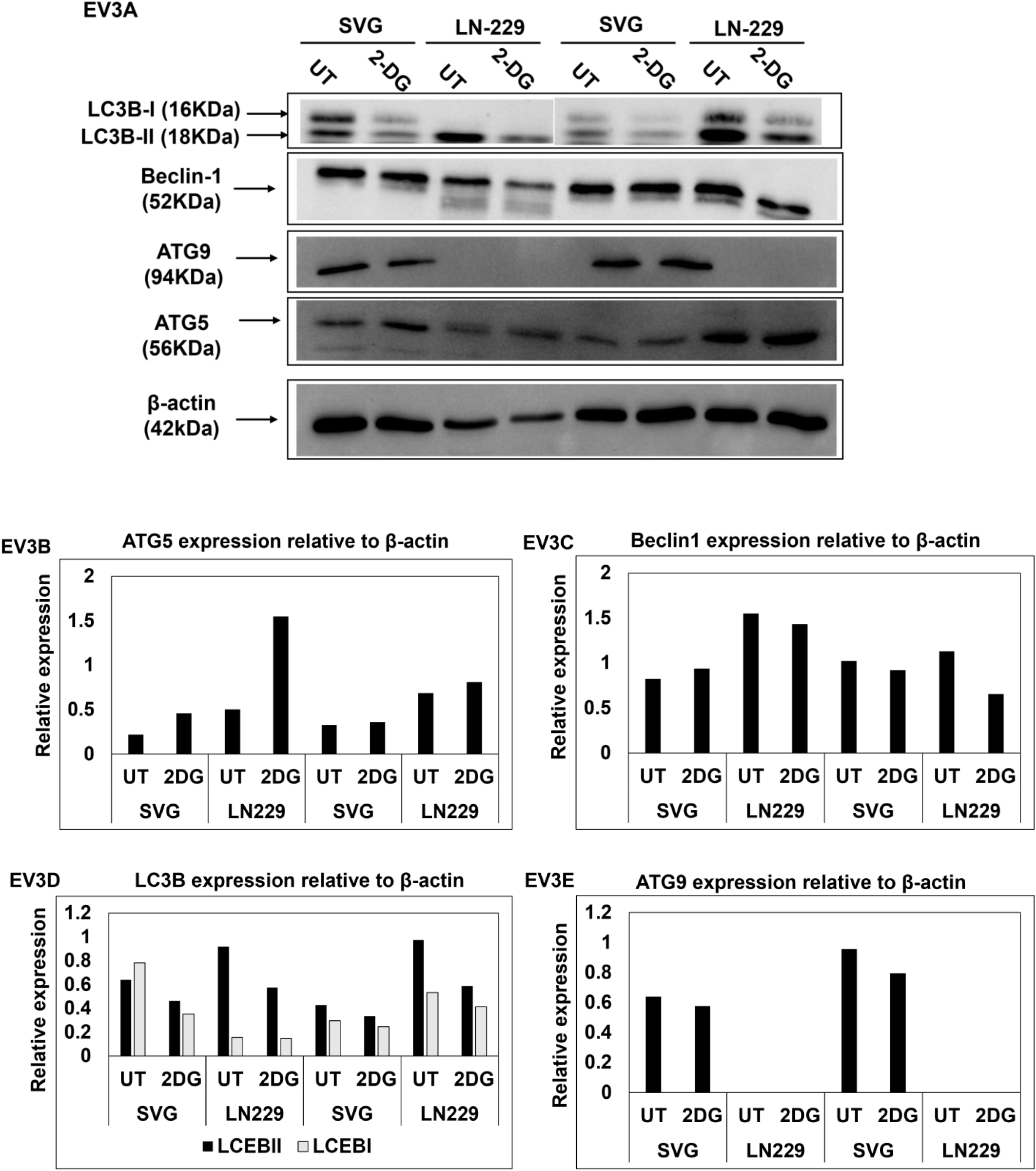
Autophagy in response to glucose starvation in GBM cell lines and astrocytes, related to Figure 6: A. LC3B, ATG5, ATG9, and Beclin-1 expression was analyzed in untreated, and 2-deoxy-glucose (2-DG) (10mM, 24hr) treated SVG and LN2-29 cells. β-actin is used as a loading control. 15µg protein sample is loaded in each well. B, C, D, E. The western blot data was quantified using densitometry (n=2)

**Fig EV4.**
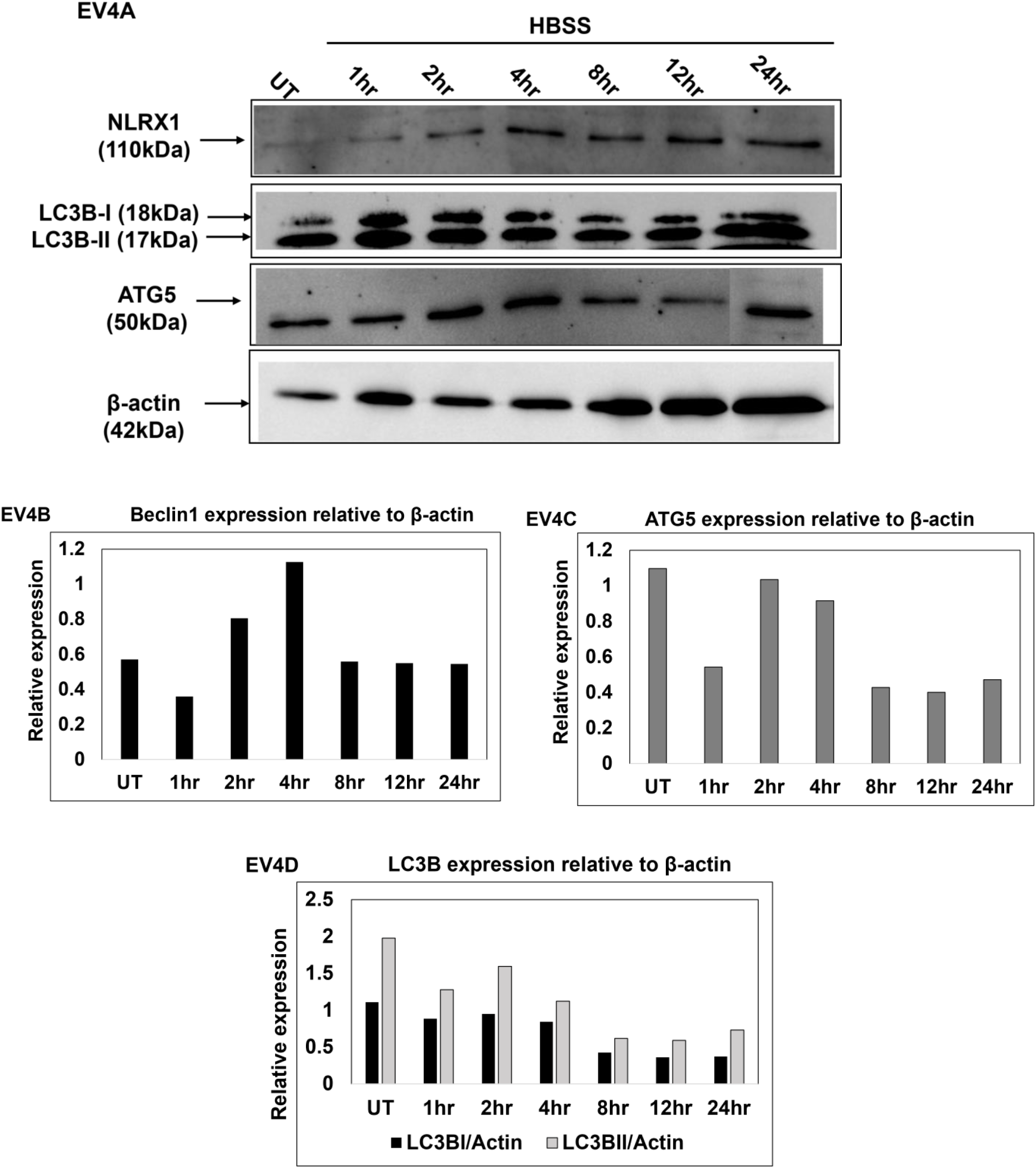
NLRX1 and autophagy markers expression in response to amino acid starvation, related to Figure 6: A. NLRX1, LC3B, and ATG5 expression were analyzed in untreated, and HBSS treated (1, 2, 4, 8, 12, 24 hours) LN-229 cells using western blot. β-actin is used as a loading control. 10µg protein sample is loaded in each well. This experiment is performed once. B, C, D. The western blot data was quantified using densitometry. The graph is representative of one experiment.

**Movie 1.**
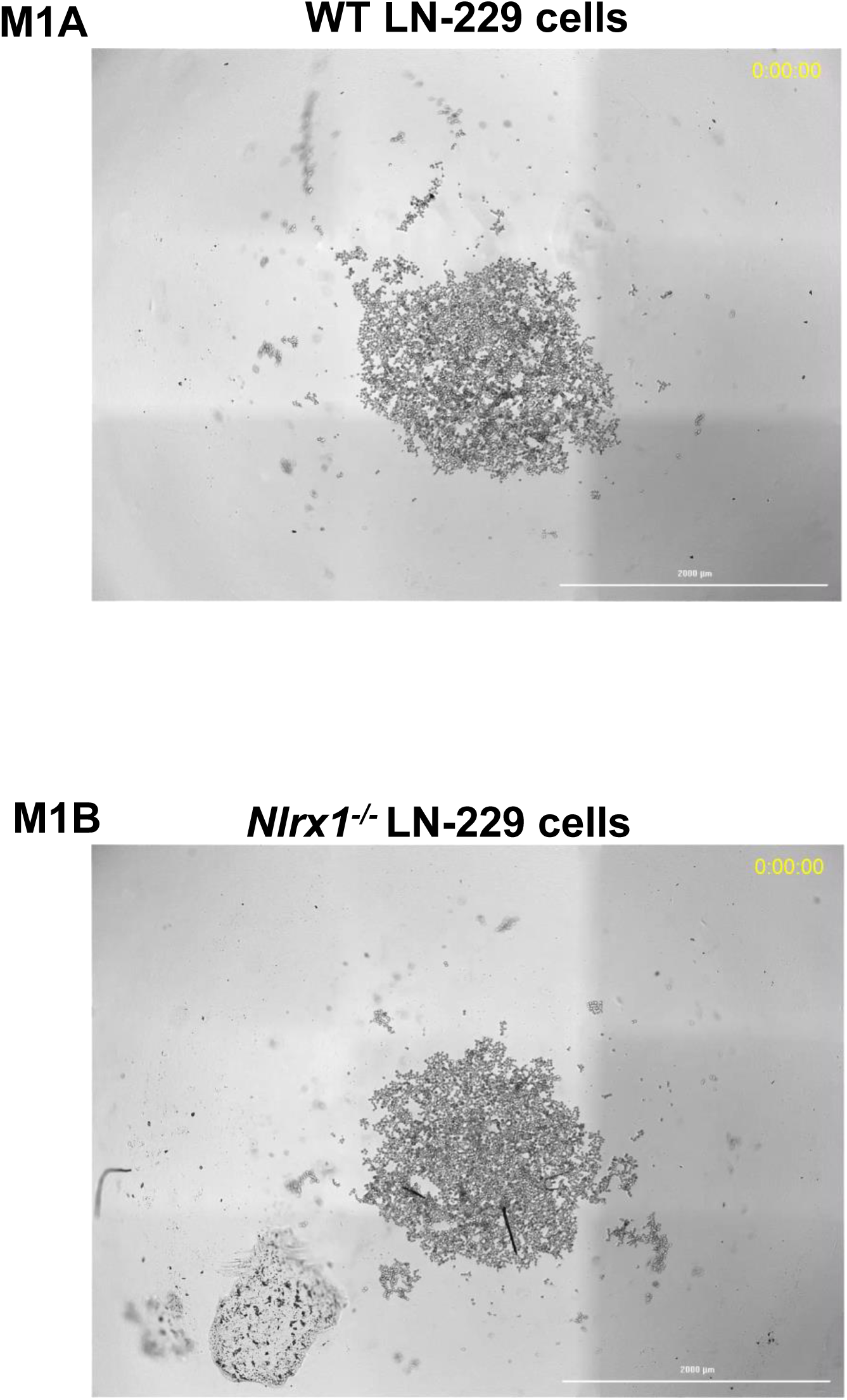
Process of spheroid generation, related to Figure 5: Movie**s** (1A) WT and (M1B) *Nlrx1^-/-^* LN-229 cells were seeded for spheroid formation, and live cell imaging was performed for 48 hours. For the creation of a movie, images were taken at intervals of 30 minutes. Movies are representative of 3 experiments. Movies were taken at a 4X objective lens, Scale bar, 2000μm.

## Notes

### Competing Interest Statement

The authors have declared no competing interest.

